# Novel AT1R-Allosteric Ligands Mask the Preeclampsia Auto-antibody Epitope and Decrease Angiotensin-induced Vasoconstriction

**DOI:** 10.1101/2021.04.30.442172

**Authors:** Khuraijam Dhanachandra Singh, Zaira P. Jara, Terri Harford, Prasenjit Prasad Saha, Triveni R. Pardhi, Russell Desnoyer, Sadashiva S. Karnik

## Abstract

Maternal blood pressure regulation by the hormone angiotensin II (AngII) sustains fetal growth through feto-placental circulation. AngII binding to orthosteric pocket in the angiotensin type 1 receptor (AT1R) induces G protein and β-arrestin signaling. AT1R blocking drugs and β-arrestin biased ligands also bind to the orthosteric pocket but evoke different inactive and active states^1–6^. AT1R-directed auto-antibodies observed in preeclampsia bound outside the orthosteric pocket to extracellular loop-2 (ECL2) of AT1R^7–9^. How auto-antibodies modulate AT1R activity causing preeclampsia pathogenesis is unknown. Here we report a druggable cryptic allosteric pocket encompassing the preeclampsia epitope on ECL2. Using structure based high-throughput small molecule screening we discovered 18 ligands specific for AT1R’s allosteric pocket. After procuring these ligands we validated inhibition of preeclampsia epitope-specific antibody binding. We characterize their inhibitory effect on antibody and AngII-signaling in cells and vasoconstriction *ex vivo*. These novel AT1R allosteric ligands, thus act as dual action negative modulators of auto-antibody action and vasoconstriction. Our study demonstrates that positive allosteric modulator action of auto-antibody causes a disease linked to AT1R. We anticipate our findings to kindle structure-based discovery of AT1R allosteric ligands for intervention in maladies such as preeclampsia^7–10^, rejection of organ transplants^11^, vasodilatory shock^12, 13^ and metabolic syndrome^14^.

## Introduction

Among disorders afflicting women, preeclampsia is the leading cause of death due to pregnancy accounting for 295,000 annual deaths globally^15^. Incidence of preeclampsia is estimated to be 1 in 25 pregnancies in the USA^16^, 1 in 10 pregnancies in Asia^17^, and one-quarter of all maternal deaths in Latin America^17^. Preeclampsia is notorious for causing mortalities of mother and fetus if the pregnancy is not prematurely terminated medically^18^. A recurring complication associated with preeclampsia is maternal hypertension causing severe intrauterine fetal growth restriction. Maintenance of maternal blood pressure and placental circulation by the renin-angiotensin system hormone, angiotensin II (AngII) through the AT1R is critical for fetal development by regulating nutrient flow and gas exchange between mother and fetus (Fig. 1a)^19^. G-protein coupled receptor (GPCR) AT 1R is the canonical AngII target that signals through Gq-mediated calcium release, β-arrestin mediated cell signaling and production of reactive oxygen species (ROS)^20^. Hyper-activation of AT1R by autoimmune antibodies might cause preeclampsia^8,10,21^. Since existing AT1R-blockers (ARBs) or Angiotensin-converting enzyme inhibitors are contraindicated in pregnancy due to their fetopathy potency, there are no treatments available for autoimmune preeclampsia^22^. Thus, intervention strategy to prevent non canonical activation of AT1R and restore normal signaling has remained an elusive goal.

**Figure 1.**
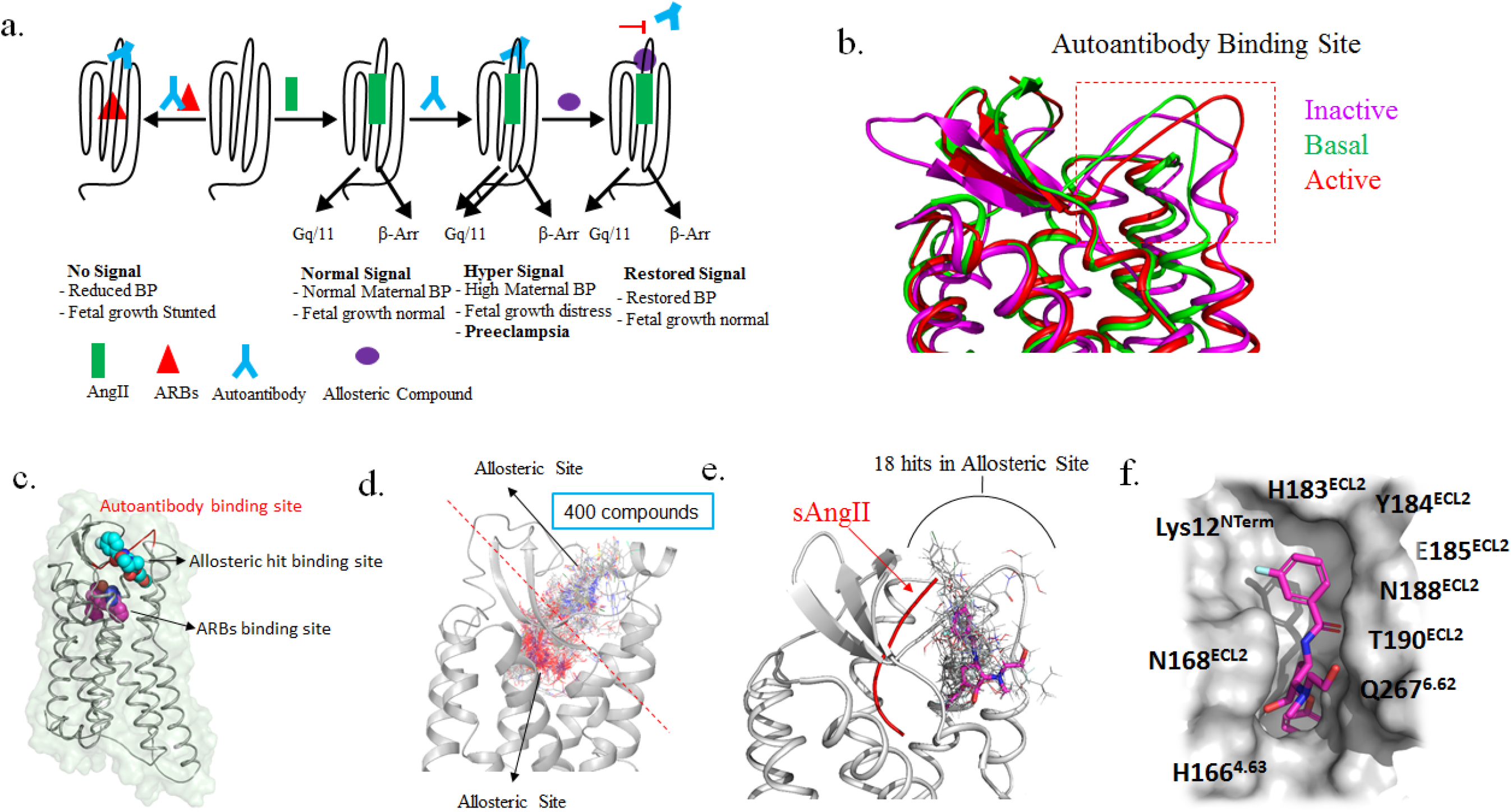
Structure based ligand discovery for the allosteric pocket of AT1R. **a**, AngII-bound AT1R signals maintain normal blood pressure and fetal growth. Autoimmune antibody binding enhance AngII signaling produces preeclampsia, maternal hypertension and fetal growth retardation. ARB binding restores maternal hypertension but crossing placental barrier exerts fetopathy. Allosteric ligand designed to inhibit autoantibody binding could restore feto-placental circulation. **b**, Conformation of ECL2-epitope in different states of AT1R. **c**, AT1R structure showing topological separation of ARB (orthosteric) and allosteric sites. **d**, HTVS result showing ligands docking to orthosteric and allosteric sites. **e,** Overlaid docking poses of 18 DCP1-compounds in relationship to bound sAngII. **f**, Typical DCP1-compound docked shown with residues in the allosteric pocket.

Canonical ligand action models conceptualize induction of active state as the physiological basis of AngII function and induction of inactive state as therapeutic basis ARBs when they bind to AT1R’s orthosteric pocket^1,3,5^. Crystal-structures of AT1R bound to agonist AngII-, β-arrestin biased agonist sAngII- and antihypertensive antagonist ARBs have validated active and inactive states of the receptor in which configuration of critical residues and motifs change^**1–6**^. The autoantibody binding ECL2 is not part of the orthosteric pocket, implying that mechanism of hyperactivation of AT1R in preeclampsia could be distinct. The epitope sequence for binding IgGs from preeclampsia patients is –<u>AFHYESQ</u>– residues 181-187 in the ECL2^23^ (Supplementary Fig. 1a), conserved in several placental mammals. Autoimmune IgGs from preeclampsia patient plasma purified and injected in mice elevate blood pressure, produce oxidative stress and renal pathogenesis in mice akin to human disease^10,24^. In previous studies we found that solvent-accessibility of –<u>CAFHYESQNST</u>– region of ECL2 is highly sensitive to different states of the AT1R and also to long-range effects of critical receptor mutations in the orthosteric pocket^25^. These findings suggested to us that dynamics of preeclampsia epitope might be influenced long-range by orthosteric ligands’ interaction with distinct pocket residues. Hence, we envisage allosteric binding of preeclampsia antibody to modulate the efficacy of orthosteric peptide agonists AngII, AngIV and non-peptide agonist L-162,313. We propose that antibody binding increases G-protein and ROS signaling. Strategically developing allosteric ligands that prevent antibody binding could be a novel treatment paradigm for autoimmune preeclampsia (Fig. 1a). Established AT1R crystal structures facilitated our high throughput virtual screening (HTVS) allowing for computational discovery and experimental validation of allosteric ligands. Allosteric ligands discovered for a few GPCRs have been approved for clinical use^26^.

### A cryptic allosteric site on AT1R

Flexibility of ECL2 became further evident from the missing electron densities for epitope residues 187-189 in the ~2.8 Å crystal structures of AT1R inactive states solved with two different antagonists (PDB ids: 4yay and 4zud)^1,2^. Flexibility might favor this region to function as a protein-protein interaction site, but the preferred conformation for autoantibody binding to AT1R is not known. Using prime suite, Schrodinger, LLC, we filled-in the missing residues in crystal structure and performed 1μs molecular dynamics simulation (MDS) as reported previously^6^. The inactive, basal and active state conformations of AT1R generated showed significant conformational changes in ECL2 (Fig. 1b). Subsequently, active state AT1R crystal structures solved also showed conformational changes in the autoantibody binding site and in the binding cleft for G-protein and β-arrestin (PDB ids: 6DO1, 6OS0) ^4,5^. Additionally, CASTp^27^ program predicted significant change of allosteric pocket shapes in different states of AT1R (Supplementary Fig. 1b-d), which makes it a cryptic allosteric site.

### Structure based HTVS

We docked ~6.8 million compounds from different libraries (Asinex, Maybridge, Chembridge, Zinc) against topographically distinct allosteric and orthosteric sites in inactive conformation of AT1R (Fig. 1c). Most of the compounds bound to the orthosteric site. Through iterative docking we confirmed that 400 compounds specifically bound to the allosteric site of AT1R (Fig. 1d). They did not bind to structurally related AT2R with different ECL2 sequence. These molecules were inspected manually for their novelty, conformational strain and lack of interaction with Trp84^2.60^, Arg167^ECL2^, Lys199^5.42^, Asp263^6.58^ and Asp281^7.32^ (superscript indicates residue position in AT1R), the orthosteric ligand binding residues (Supplementary Fig. 1a). Following this, 400 compounds were subjected to Glide XP docking in all three states of AT1R resulted in 197 compounds capable of stable binding to the allosteric pocket in inactive, basal and active states. To ensure diversity of final compounds, we performed clustering analysis based on their structural similarity using Canvas Similarity and Clustering module of Schrodinger. The 197 compounds were grouped into seven clusters (Supplementary Fig. 2). A representative compound with the best docking score was selected from each cluster. A structural similarity search performed using Sci-finder (https://scifinder-n.cas.org) yielded eleven more compounds. The compounds were named as DCP1-1 through DCP1-18 in order of their discovery. Chemically seven belonged to cyclopentane derivatives, two to pyrrolidine derivatives and six to piperidine derivatives. DCP1-1, −2 and −5 did not belong to above groups and have different structures (Supplementary Fig. 3). Next we performed Glide Induced Fit Docking (IFD) in which the receptor side chains are flexible to give an accurate binding pose (Supplementary Fig 4 and Supplementary Table 1). Eighteen compounds showed docking score of less than −7 and overlapping poses for the allosteric pocket (Fig. 1d). In addition to ECL2, residues from transmembrane helices, TM4, TM6 and N-terminus contribute in the allosteric pocket (Fig. 1e). Thirteen out of 18 compounds interact with His166^ECL2^, 16 with His183^ECL2^ and 12 with T190^ECL2^. These 18 compounds were acquired (from Asinex, Winston-Salem, North Carolina and Aurora Fine Chemicals LLC, San Diego) for experimental validation studies.

### Inhibition of antibody binding to preeclampsia epitope

Purified rabbit polyclonal antibody (IgG) for ECL2-sequence, –<u>CIENTNITVSAFHYESQNS</u>– was used as a surrogate for preeclampsia auto-antibody in validating the DCP1-compounds (Fig. 2). Concentration dependent binding of this IgG to HEK293 cells permanently expressing AT1R (HEK-AT1R)^1^ was inhibited specifically by antigenic peptide but not by orthosteric ligands AngII and Olmesartan (Supplementary Fig. 5a-c). The IgG potentiated AT1R signaling by agonists, AngIV, AngII and L-162,313 (Fig. 2c). A similar anti-AT1R antibody which potentiated AngII signaling was previously reported^28^. All 18 compounds inhibited IgG binding to HEK-AT1R cell-surface (Fig. 2a). Relative to the ECL2-peptide, inhibition of IgG binding by several DCP1-compounds was significantly more (p<0.001). Specifically, IC_50_ values of DCP1-3 and DCP1-16 were 2.9 and 0.6 nM respectively (Fig. 2b). Inhibition of IgG binding by DCP1-3 and DCP1-16 led to inhibition of the antibody-effect on agonist-induced calcium response (Fig. 2c, d) and ROS production (Fig. 2e). These data reveal that DCP1-3 and DCP1-16 effectively block antibody binding to AT1R as well as the agonist potentiating effects of the IgG.

**Figure 2.**
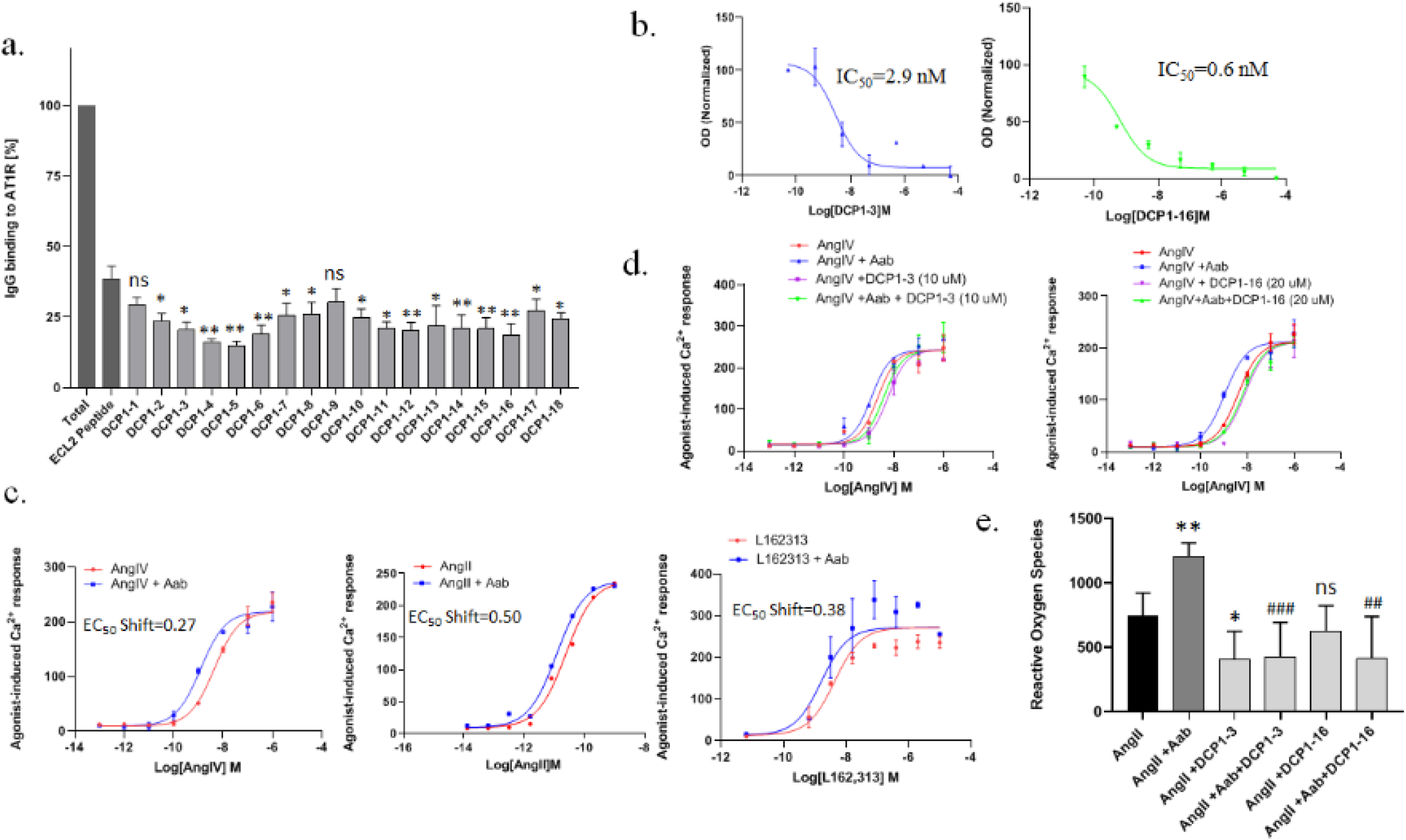
Allosteric ligand effects on IgG binding to the preeclampsia epitope. **a**, Binding of IgG to HEK-AT1R cells without and with addition of allosteric ligands. Significance of inhibition compared to the antigenic peptide is shown **p<0.001, *p<0.05 and ns, not significant. **b**, Potency of IgG binding inhibition by DCP1-3 and DCP1-16. **c**, IgG binding positively modulates potency of three AT1R agonists. **d**, Reversal of IgG-induced agonist potency-shift by DCP1-3 and DCP1-16. **e**, IgG potentiated AngII-induced ROS production inhibited by DCP1-3 and DCP1-16. * p<0.05 change compared to AngII, ## p<0.05 and ### p<0.001 change compared to IgG+AngII and ns, not significant.

### Allosteric property of compounds

Apart from inhibiting IgG effects on AT1R, the DCP1-compounds modulate affinity and efficacy of orthosteric agonists. Varying the concentrations of DCP1-compounds did not alter equilibrium binding of ^125^I-[Sar^1^Ile^8^]AngII and did not elicit agonist-like calcium response (Supplementary Fig. 6a, b). However, concentration dependent binding of agonist ^125^I-AngIV is reduced in the presence of 50 μM DCP1-3 and DCP1-16 (Supplementary Fig.6c), which also reduced AngIV signaling responses (Fig. 3), indicating that their negative allosteric modulator (NAM) activity manifested only in the presence of orthosteric agonists. Binding of agonist ^125^I-AngII was not significantly reduced by DCP1-compounds.

**Figure 3.**
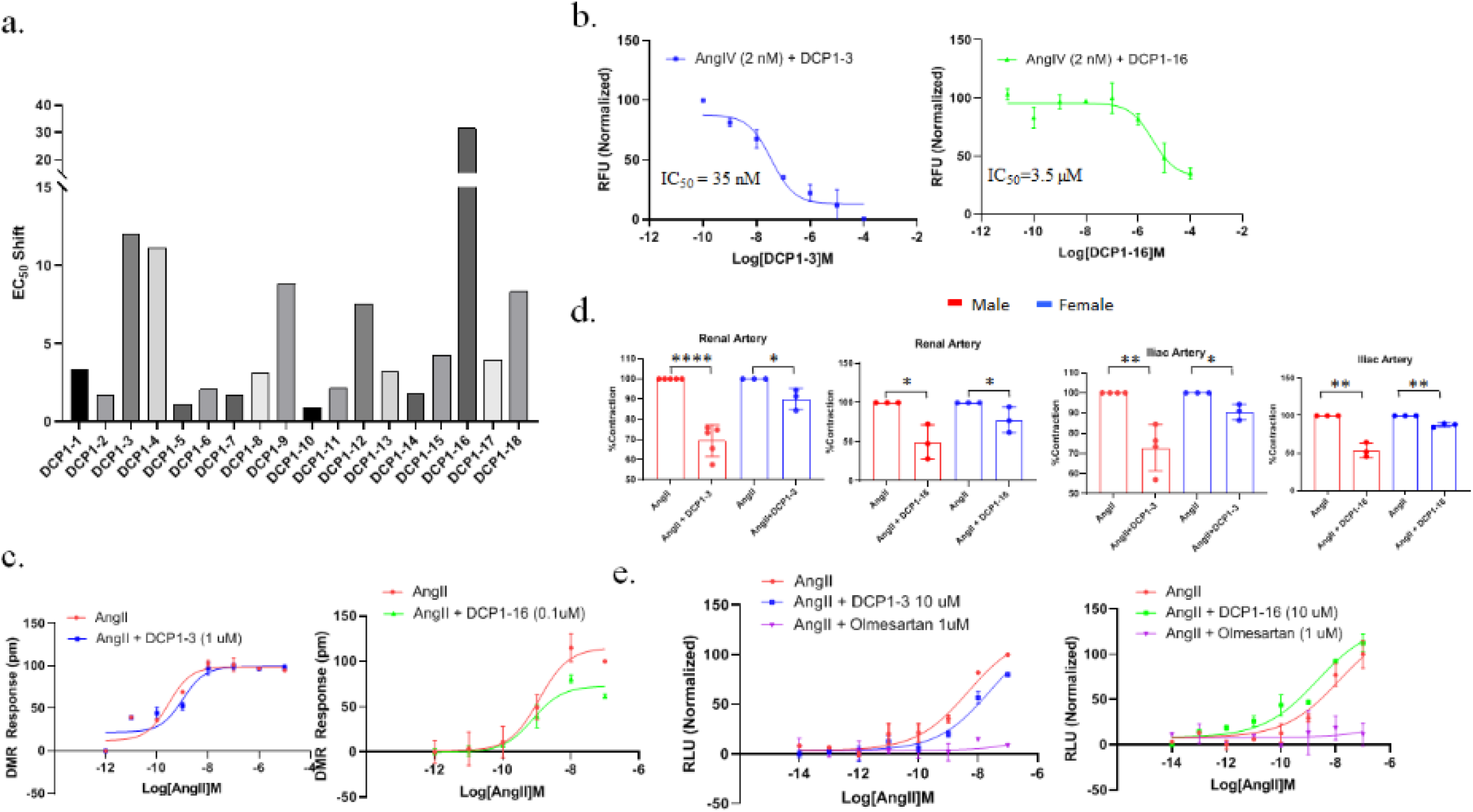
Efficacy modulation of orthosteric agonist effects: **a**, Fold decrease of EC_50_ for AngIV-induced Ca^2+^ response in presence of 30 μM of indicated allosteric ligand. A shift of >10-fold was used as cut-off for selecting DCP1-3 and DCP1-16 for additional studies. **b**, NAM potency of DCP1-3 and DCP1-16 on AngIV at EC_50_ concentration. **c**, NAM effect of DCP1-3 and DCP1-16 on AngII-induced DMR dose response. Data are mean ± SD (n=3 measurements). **d**, Effects 30 μM DCP1-3 and DCP1-16 on 200 nM AngII-induced vasoconstriction *ex vivo*. Each bar represents average ± S.D of response readings in the number of mice indicated in filled circles on each bar. Male and female mice were used because vascular response to AngII is influenced by gender. *** indicates p<0.001, ** indicates p<0.05, * indicates p=0.05. **e**, β-arrestin recruitment induced by various ligand mixtures assayed by TANGO assay. Data are mean ± SD (n = 3 measurements).

We monitored DCP1-compound effects on agonist-dependent AT1R signaling at several levels, intracellular Ca^2+^ release which is linked to ROS production and vasoconstriction as well as β-arrestin recruitment and global cellular changes (Fig. 3). The EC_50_ of AngIV dose-response curves increased in the presence of all DCP1-compounds. Increase was >10-fold for DCP1-3, DCP1-4 and DCP1-16 (Fig. 3a). The AngIV response at EC_50_ dose was systematically inhibited by increasing concentrations of DCP1-3 and DCP1-16 (Fig 3b and Supplementary Fig. 7). Negative co-operativity suggests reciprocal regulation of allosteric and orthosteric pocket conformations, typically manifested as affinity and efficacy modulation of orthosteric agonists^29^. As shown affinity for ^125^I-AngIV is reduced in the presence of DCP1-3 and DCP1-16 (Supplementary Fig. 6c). Consequently, 30 μM DCP1-3 and DCP1-16 caused 12- and 31-fold decrease of agonist efficacy (Fig. 3a and Supplementary Fig. 7a-b). NAM potency of DCP1-3 was estimated as IC_50_= 35 nM at EC_50_ and no NAM effect was observed at EC_80_ of AngIV. DCP1-16 NAM effect at the EC_50_ concentration of AngIV was stronger, IC_50_ = 3.5 nM (Fig. 3b).

NAM effect of DCP1-3 and DCP1-16 on AngII-simulated Ca^2+^ response was less pronounced, and more similar to a neutral allosteric ligand (NAL) effect. Such modulators are also important pharmacological tools. To assess effect of DCP1-3 and DCP1-16 on additional AngII signals we used the label free dynamic mass redistribution (DMR) assay and *ex vivo* vasoconstriction assay. DMR is a sensitive measure of global cellular changes brought about by receptor signaling without the use of chemical dyes or labels to monitor cellular events. DMR signal changes were AngII dose dependent and the signal was completely inhibited by ARB treatment (Supplementary Fig. 8a). In presence of DCP1-3 and DCP1-16 the DMR-response curves right-shifted (Fig. 3c), demonstrating NAM effect on AngII-mediated cellular signaling by the AT1R.

We next examined efficacy of these compounds in an *ex vivo* model: constriction of live renal and iliac arterial segment explants from mouse in a myograph chamber (Supplementary Fig. 9a). To ensure an intact endothelium and vessel behavior, responses to KPSS, phenylephrine and acetyl-choline were initially assessed. Responsive arterial segments were then exposed to 200 nM AngII, or 200 nM AngII+DCP1-3 (30 μM) and 200 nM AngII+DCP1-16 (30 μM) and vasoconstriction was monitored (Supplementary Fig. 9b). We found significant reduction (p<0.05) in vasoconstriction by AngII when treated with DCP1-3 and DCP1-16, in both renal and iliac arteries from male and female mice (Fig. 3d). The inhibitory effect was reversed by washing the vessels and re-stimulating with AngII reproduced contraction (Supplementary Fig. 9b). This *ex vivo* experiment further exemplifies the NAM properties of DCP1-3 and DCP1-16.

Induction of signal bias by allosteric ligands is an emerging concept of significant interest^30–31^. In order to determine potential β-arrestin bias exerted by DCP1-3 and DCP1-16, we utilized the TANGO assay^32^. Cells were exposed to either AngII alone or AngII with either DCP1-3 or DCP1-16 (10uM). Addition of DCP1-3 resulted in a reduced level of β-arrestin recruitment signal compared to AngII alone. However, in DCP1-16 treated cells β-arrestin recruitment signal increased (Fig. 3e). This result revealed that DCP1-3 is a NAM for both G-protein and β-arrestin signals but DCP1-16 is a NAM for G-protein- and PAM for β-arrestin-recruitment.

### Mechanism of the allosteric effect

Molecular architecture of DCP1-3 and DCP1-16 includes same functional groups at position 2 and 3 of a cyclopentane scaffold. Comparison between their binding modes (Fig. 4a) show that both interact with the preeclampsia epitope residues His183^ECL2^, Tyr184^ECL2^ and neighboring residues Glu185^ECL2^, Thr190^ECL2^, and Leu191^ECL2^. This can account for masking the preeclampsia epitope from IgG binding as observed in Fig. 2a. In addition, both interact with Pro162^4.59^, His166^4.63^, Asn168^ECL2^ and Phe170^ECL2^ which can account for their NAM potency seen in Fig. 3a. The oxygen in the tetrahydropyran ring of DCP1-3 interacts with Tyr184^ECL2^. The methoxy oxygen of DCP1-16 interacts with Gln267^6.67^. Each of these is a unique interaction that could account for pharmacological differences observed between DCP1-16 and DCP1-3 in Fig. 3e.

**Figure 4.**
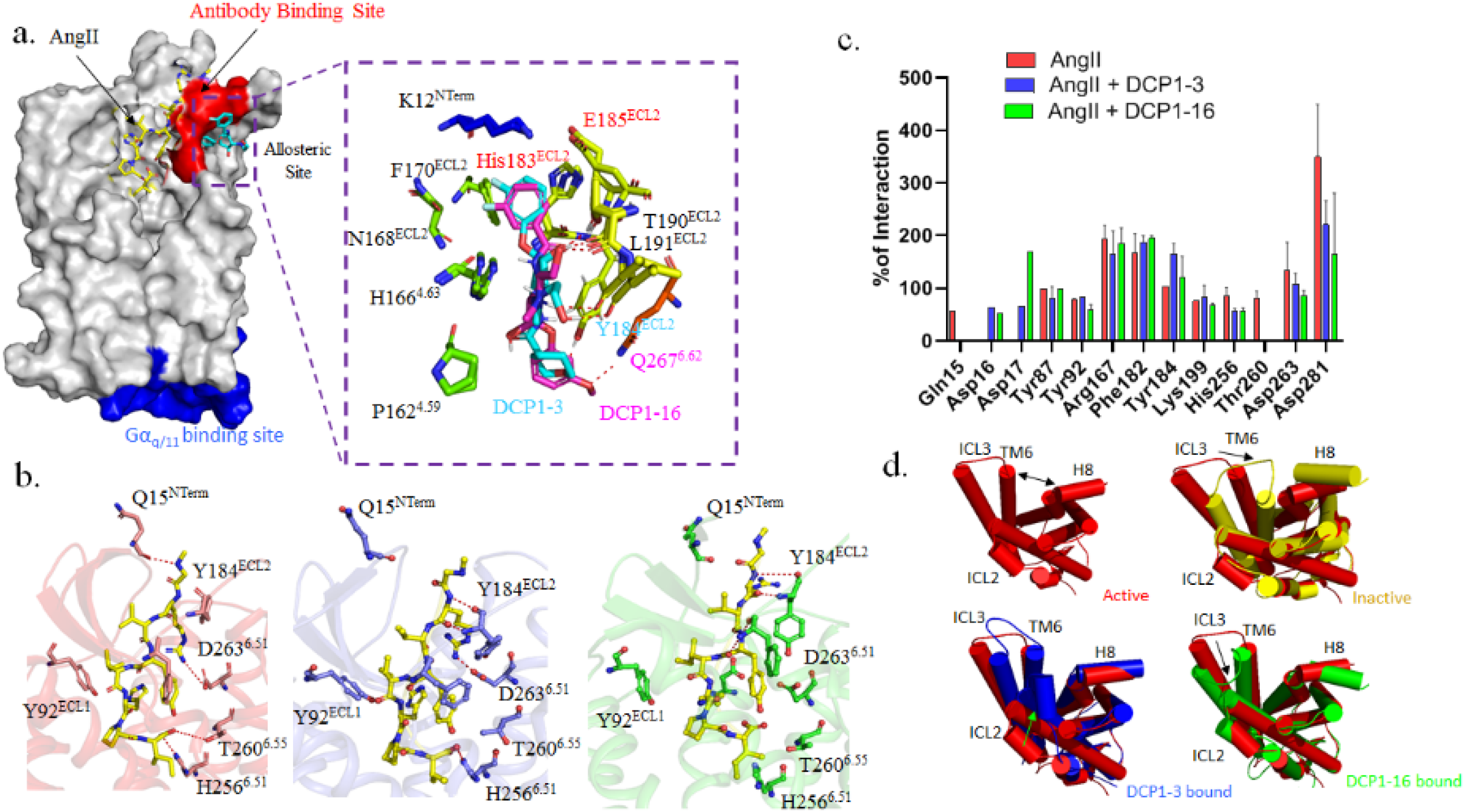
Effects of bound allosteric ligands on AT1R structure and activation state. **a**, AT1R with AngII-bound to orthosteric pocket and DCP1-3 bound to allosteric pocket. Enlarged inset of allosteric pocket shows overlay of DCP1-3 and DCP1-16 poses. Sidechains of interacting residues change orientation to accommodate each compound structure. Residues within the epitope for IgG are shown in red. Residue interaction unique to each DCP1-compound is shown in same color as the compound. **b**, Interactions of AT1R residues with orthosteric peptide ligand sAngII (left) and in the presence of DCP1-3 (middle) and DCP1-16 (right). Note changes in distance of interaction of each orthosteric sidechain in presence of DCP1-3 and DCP1-16, which may reduce affinity and efficacy of the orthosteric agonists, i.e. NAM. **c**, Change of % interaction of critical orthosteric pocket residues induced by DCP1-3 (blue) and DCP1-16 (green) compared to % interaction in their absence (red). **d**, Changes in the orientation of ICL2, ICL3, TM6 and H8 depicted in the cytoplasmic view. Changes induced by DCP1-3 and DCP1-16 compared to active-state of AT1R affect the overall size of binding cleft for G-protein and β-arrestin engagement.

To identify structural changes induced when a NAM binds to AngII engaged AT1R (Fig. 4a), we compared the structures of different complexes at the end of 1000 ns MDS. The overall structures of the DCP1-3+sAngII+AT1R and DCP1-16+sAngII+AT1R were similar to sAngII+AT1R. However, structural differences were observed in ECL2, the orthosteric pocket and the cytoplasmic cleft when allosteric ligands bind. ECL2 stability increases; consequently the reduced entropic state of ECL2 may prevent preeclampsia antibody binding. In the orthosteric pocket the interactions of sAngII with receptor decreased as observed from MM/GBSA free energy decrease upon binding DCP1-3 for DCP1-16 (Supplementary Tables 1and 2). Overall percentage of interactions of orthosteric pocket residues, His256^6.51^, Thr260^6.55^, Asp263^6.58^, and Asp281^7.32^ was reduced (Fig. 4b, c). These are critical residues for agonist-induced activation of AT1R^5^. These changes may account for reduction of agonist binding and decrease of signaling efficacy shown in Fig. 3. In the cytoplasmic region, ICL2, ICL3, TM6 and H8 of AT1R show different motions in response to DCP1-3 and DCP1-16 (Fig. 4d). As a result, the size of the cleft for binding G-protein or β-arrestin decreases upon binding DCP1-3. However, there was no reduction in the size of the cleft upon DCP1-16 binding when compared to the β-arrestin biased sAngII bound AT1R structure^4^. This difference explains β-arrestin biased property of DCP1-16 observed in Fig. 3e. Together, these structural differences induced can be signature changes for transition of AT1R from the IgG-bound overactive disease producing state to the NAM-bound state that can account for near-normal AngII signals as depicted in Fig. 1a.

In summary, the preeclampsia epitope is part of a cryptic allosteric site of AT1R. For the first time we discovered non-peptide small molecules. When bound they mask the preeclampsia epitope from IgG binding, but are weak NAMs for AngII signaling. This is beneficial to avoid potential harm to fetus due to complete inhibition of AngII signals as it happens in the case of ARBs. Developing autoimmune therapy through designed drugs may be widely beneficial. For instance, AT1R autoantibodies cause a variety of pathologies including renal graft rejection^10^ and transplantation failure of heart^11^, hand^33^, liver^34^, lung^35^ and stem cells^36–37^ for which there are no current treatments. This is also the first report of allosteric ligands of AT1R. Through structure-based HTVS we recovered small molecule allosteric hits with cyclopentane, pyrolidine and piperidine scaffolds. This new knowledge provides the foundation for future design of novel AT1R allosteric ligands as potential lead candidates for detailed biological evaluation as potential NAM, PAM and biased ligands. AT1R PAM ligands may be especially useful in treatment of hypotensive disorders^12–13^. Increasing the potency of allosteric ligands for specialized pharmacological modulation through rational drug design approaches could lead to a new generation of AT1R targeted therapeutics.

## Methods

### Protein Preparation

The starting coordinates of AT1R [PDB IDs: 4ZUD]^1^ were retrieved from Protein Data Bank (www.rcsb.org) and further modified for Glide docking calculations (Glide, Schrodinger, LLC, NY, USA)^3,37^. This work was completed already before the sAngII or AngII or TRV bound AT1R structures were solved (PDB ids: 6do1, 6os0, 6os1, 6os2)^4,5^. For the calculations, energy minimization was applied to the protein using the Protein Preparation Wizard by applying an OPLS3 (Optimized Potentials for Liquid Simulations) force field^39^. Missing segments of loops were filled-in using Prime, Schrodinger, LLC, during the protein preparation^40^. Progressively weaker restraints applied to only the non-hydrogen atoms and structure refinement was done based on the default protocol of Schrodinger, LLC, NY, USA, as Glide uses the full OPLS3 force field at an intermediate docking stage and is claimed to be more sensitive to geometrical details than other docking tools. The most likely positions of hydroxyl and thiol hydrogen atoms, protonation states, and tautomers of His residues, and Chi “flip” assignments for Asn, Gln, and His residues were selected. Finally, energy minimizations of protein were performed until the average root-mean-square deviation of the non-hydrogen atoms reached 0.3 Å. Inactive state structure of AT1R [PDB ID: 4ZUD], basal state and AngII bound or active state structure were generated from our molecular dynamics simulation study (MDS)^3,6^. Five average structures of basal state and active state structure were generated from our MDS. These structures were used for our initial structure based screening of compound library.

### Compound database

The compounds were collected from Zinc, Asinex, Maybridge, Chembridge database (~6.7 million). Lipinski-filtered compounds were cleaned using LigPrep^41^, with expansion of stereo centers whose chirality was unspecified. This small molecules collection contains a high percentage of drug like compounds.

### High Throughput Virtual Screening (HTVS)

Glide implements a highly efficient virtual screening workflow (VSW) that is able to rank compounds through fast high-throughput virtual screening and then docking calculations of higher accuracy using standard and extra precision. For the allosteric site, the Center of Mass (COM) position of the residues are 181, 182, 183, 184, 185, 186, 187 and was set as the grid center. The inner and outer box values were set to 10Å and 25Å, respectively^42^.

The compounds were subjected to Glide based three-tiered docking strategy in which all the compounds were docked by three stages of the docking protocol, High Throughput Virtual Screening (HTVS), Standard Precision (SP) and Extra precision (XP). First stage of HTVS docking screens the ligands that are retrieved and all the screened compounds are passed on to the second stage of SP docking. The SP resultant compounds were then docked using more accurate and computationally intensive XP mode. Based on the glide score and glide energy, the protocol gives the leads in XP descriptor. Glide module of the XP visualizer analyses the specific interactions. Glide includes ligand-protein interaction energies, hydrophobic interactions, hydrogen bonds, internal energy, π-π stacking interactions and root mean square deviation (RMSD) and de-solvation^42–45^.

### Induced Fit Docking (IFD)

Induced fit docking (IFD) (Schrodinger, LLC, NY, USA) was used to dock allosteric hits to our proposed allosteric site. First, the ligand was docked into a rigid receptor model with scaled down vdW radii. A vdW scaling of 0.5 was used for both the protein and ligand nonpolar atoms. A constrained energy minimization was carried out on the protein structure, keeping it close to the original crystal structure while removing bad steric contacts. Energy minimization was carried out using the OPLS3 force field with implicit solvation model. Glide XP mode was used for the initial docking and ligand poses were retained for protein structural refinements. Prime (Schrodinger, LLC, NY, USA) was then used to generate the induced-fit protein-ligand complexes (Prime, Schrodinger, LLC, NY, USA). Each of the structures from the previous step were subjected to side-chain and backbone refinements. All residues with at least one atom located within 4.0 Å of each corresponding ligand pose were included in the Prime refinement. The refined complexes were ranked by Prime energy, and the 20 receptor structures within 30 kcal/mol of the minimum energy structure were passed through for a final round of Glide docking and scoring. In the final step, each ligand was re-docked into the top 20 refined structures using Glide XP^45,46^.

### MM/GBSA

Prime/MM-GBSA was used to predict the free energy of binding between the receptor and the AngII. The binding free energy (ΔG_bind_) was calculated using Prime^**6**^. The binding free energy (G_bind_) is then estimated using the following equation:

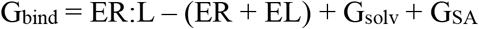

where ER:L is energy of the complex, ER + EL is the sum of the energies of the ligand and the apo protein, using the OPLS3 force field, and ΔG_solv_ (ΔG_SA_) is the difference between GBSA solvation energy (surface area energy) of complex and sum of the corresponding energies for the ligand and apoprotein^47–48^.

### MD Simulation

Molecular dynamics (MD) simulations were carried out for AT1R-apo and AT1R-AngII complexes using Desmond MD code and the OPLS3 force field^38^ for minimization of the system^6^. Also, we ran the MD simulation for 1 μS for each of the complexes (AT1R-sAngII, AT1R-sAngII-DCP1-3, AT1R-sAngII-DCP1-16). Using the Desmond system builder, a 10 Å buffered orthorhombic system with periodic boundary conditions was constructed using a POPC lipid membrane and an SPC explicit water solvent. The overall charge was neutralized by 0.15 mol/ L NaCl. The simulations were performed in the NPT ensemble with number of elements, pressure, and temperature controlled. The temperature of 300 K and pressure of 1.013 bar were kept constant by coupling the system to a Berendsen thermostat and barostat, which are normal temperature and pressure mimicking the real life environment. An integration step of 2.0 was used, Coulombic interactions were calculated using a cutoff radius of 9.0 Å and long-range electrostatic interactions were calculated using the smooth particle mesh Ewald method. Before each MD simulation, a default Desmond membrane protein relaxation protocol was applied^49–52^. All the figures were generated using PyMol (The PyMOL Molecular Graphics System, Version 2.0 Schrödinger, LLC).

### ^125^I-Ang peptide Specific Binding Analysis

Ligand binding was analyzed using membranes prepared from HEK293T cells expressing wild-type HA-AT1R as described earlier^2,24^ and suspended in membrane binding buffer (140 mM NaCl, 5.4 mM KCl, 1 mM EDTA, 0.006% bovine serum albumin, 25 mM HEPES, pH 7.4). A 10 μg of homogeneous cell membrane was used per well. Competition binding assays were performed under equilibrium conditions, with 2 nM radioligand (^125^I-AngII, ^125^I-[Sar^1^, Ile^8^]AngII and ^125^I-AngIV are purchased from Dr. Robert Speth, NOVA University, FL) and concentrations of the competing ligand ranging between 0.04 and 1000 nM. Binding reaction was terminated by filtering the binding mixture through Whatman GF/C glass fiber filters, washed with buffer (20 mM sodium phosphate, 100 mM NaCl, 10 mM MgCl2, 1 mM EGTA, pH 7.2). The bound ligand concentration was determined as the counts/min (MicroBeta2 Plate Counter, PerkinElmer Life Sciences). Nonspecific binding was measured in the presence of 105 M ^127^I-AngII (Bachem). 3-5 independent experiments, each done in triplicate, were performed and the resulting values were pooled to a mean curve which is displayed. The binding kinetics were analyzed by the nonlinear curve-fitting program GraphPad Prism 8. The means ± SE for the IC_50_ values were calculated^2^.

### Effect on saturation binding of peptide agonists

Saturation binding assays with ^125^I-AngIV were performed under equilibrium conditions, with ^125^I-AngII or ^125^I-AngIV concentrations ranging between 1.6 and 27 nM (specific activity, 16 Ci/mmol) as duplicates in 96-well plates for 1 hr at room temperature in the presence of 50 μM DCP1-3 and DCP1-16 compounds. The binding kinetics were analyzed by nonlinear curve-fitting program GraphPad Prism 8, which yielded the mean ± SE for the Kd and Bmax values^1,2^.

### ECL2 antibody production in rabbit and affinity purification/characterization of IgG specificity

The selected peptide antigen -CIENTNITVSAFHYESQNS- was conjugated to Keyhole Limpet Hemocyanin (KLH). The conjugated peptide was injected into 2 rabbits at 2-3 week intervals 5 times. Blood was collected 7 days after the 3^rd^ and 4^th^ injection. A terminal bleed was done after the 5^th^ injection. Specific antibody titers for each bleed were measured by ELISA using free peptide -CIENTNITVSAFHYESQNS- coated to the plate (4ug/ml, 50ul/well, 96 well, in carbonate buffer overnight at 4 degrees). The antigenic peptide -CIENTNITVSAFHYESQNS- was linked to a resin using Pierce’s Sulfo-Link Immobilization Kit for peptides (Pierce, Catalog # 44999). Peptide-specific antibody from 20 ml of serum was purified using a 2 ml antigenic peptide-affinity column and used in the ELISA and Ca^2+^ mobilization assays described in the methods.

### ELISA assay

HA-AT1R expressing cells were seeded at a density of 600,000 cells/well in poly-l-lysine coated 24-well clear bottom plates. After overnight incubation, plates are washed with WB1 (1% BSA in HBSS, 500ml + 5g BSA). Cells were then fixed with 4% paraformaldehyde at room temperature for 15 minutes and washed 2x with WB1. Cells were treated with DCP1-compounds for 1 hour at room temperature. After 1 hour, AT1R ECL2 autoantibody (Aab) was added (1:1000) to all the wells treated with DCP1 compounds and incubated for an additional 1 hour. Plates are then washed with WB1, rabbit secondary antibody is added to all wells and plate was incubated at room temperature for 1 hour. Plates are then washed with WB2 (0.5% BSA in HBSS, 500ml + 2.5g BSA). 200 μl OPD (Phosphate-Citrate Buffer with Sodium Perborate + o-Phenylenediamine dihydrochloride) solution is added to each well and color development is allowed to proceed for 15 minutes. 3N HCl is added to stop the reaction and absorbance is read at 492 nm on the Flexstation 2, Molecular Devices. The percentage of inhibition is presented for each compound. Data were normalized and analyzed using the sigmoidal dose-response function built into GraphPad Prism 8.0.

### Comparative Agonism by Intracellular Calcium Levels

Calcium levels were measured using Fluorescent Imaging Plate Reader (FLIPR) calcium 5 assay kit (Molecular Devices) as described previously^1^. HA-AT1R expressing cells were seeded at a density of 100,000 cells/well in a 96-well clear bottom black plate. Following serum starvation, calcium-sensitive dye was added to the cells. The Flexstation instrument was programmed in FLEX mode to add ligands (0.04 and 1000 nM concentration) to the cells and to monitor the fluorescence before and after adding the ligands. The dose-response curves were plotted and percent agonism was calculated assuming 100% stimulation by AngII. Changes in intracellular calcium were recorded by measuring ΔF/F (max-min), and represented as relative fluorescence units (RFU). Data were analysed using the sigmoidal dose–response function built into GraphPad Prism 8.0.

#### Vascular reactivity

Vascular contraction studies were conducted following the National Institutes of Health’s guidelines for care and use of animals and with approved mouse protocols from the institutional animal care and use committees. On the day of experimentation, C57BL/6 male and female mice were euthanized by isoflurane overdose followed by cervical dislocation. 1.5-2 mm renal and iliac arteries were removed, transferred to a dish containing physiological saline solution (PSS: 130 mM NaCl, 4.7 mM KCl, 1.18 mM KH2PO_4_, 1.17 mM MgSO_4_, 14.9 mM NaHCO3, 5.5 mM Dextrose, 26 μM CaNa_2_ Versenate, 1.6 mM CaCl_2_) and perivascular fat was removed. Vessels were then mounted in multi-wire myograph system (Danish Myo Technologies (DMT) model 620N using 40 μm stainless steel wires. Vessels were incubated in warm PSS for 60 minutes with 5% CO_2_ - 95% O_2_ and aerated at a constant rate. Normalized passive resting tension was determined and vessels were allowed to equilibrate for 20 minutes. The integrity of the vessels were assessed by a) voltage-dependent contraction which was induced by receptor-independent depolarization of the vascular smooth mucle cell membrane with elevated K^+^. PSS was replaced with 60 mM KCl isotonic depolarizing buffer (KPSS): (74.7 mM NaCl, 60 mM KCl, 1.18 mM KH_2_PO_4_, 1.17 mM MgSO_4_, 14.9 mM NaHCO_3_, 5.5 mM dextrose, 26 μM CaNa_2_ Versenate, 1.6 mM CaCl_2_) causing contraction, after reaching contraction plateau, KPSS was washed out and replaced with PSS. b) receptor-dependent contraction and relaxation which were induced by α_1_-adrenergic and muscarinic stimulation with phenylephrine and methylcholine, respectively. 200 nM phenylephrine was added into the chamber containing PSS, further the contracted vessels were then relaxed by adding 1 μM methylcholine. Vessels that passed the integrity test were used to assess AngII-dependent contraction. Responses were recorded by using computerized data acquisition and recording software LabChart 8 (AD Instruments, Colorado Springs CO).

#### AngII-dependent contraction

Following vessel response and integrity testing, AngII stimulation over a 3-hour period caused contraction in renal and iliac arteries from both male and female. AngII-dependent contraction levels were lower in females than males, perhaps due to higher expression of AT2R increasing steady-state levels of NO exclusively in this gender opposing AT1R action^53^. To overcome this gender effect, AngII contraction in female arteries was performed in the presence of 100umol/L of N(gamma)-nitro-L-arginine methyl ester (L-NAME), a nitric oxide synthase inhibitor. To test the efficacy of DCP1-compounds, vessels were contracted with 2 nM AngII in PSS. Once the contraction returned to baseline, AngII was washed out. After one hour a second stimulation with AngII in combination with 30 μM DCP1-3 or DCP1-16 was performed. Once the stimulation returned to baseline, AngII +DCP1-3 or DCP1-16 was washed out. One hour after a third stimulation with AngII alone was performed to ascertain preservation of the vessel through the experiment. An hour after AngII washed out, the final treatment was with AngII+ARB to demonstrate AT1R-selectivity.

Sample sizes (number of animals) were not predetermined by a statistical method and animals were assigned to groups randomly. Predefined exclusion criteria used was lack of AngII-induced contraction, following assessment of KPSS, phenylephrine and acetylcholine response in vessels obtained from both genders. Statistical analyses were performed after first assessing the normal distributions of data sets and assessment of variances. In Fig. 4d, an asterisks, * P < 0.05, ** P < 0.005, *** P < 0.0005 and **** P < 0.0001 each indicating significance of difference between AngII and AngII+DCP1- (calculated using unpaired t-test).

#### Dynamic mass redistribution (DMR) studies on HEK-AT1R cells

DMR experiments was performed on HEK-AT1R stable cell line by seeding the cells into EPIC-corning fibronectin coated 96-well DMR microplates (Cat. 5082-Corning) with a density of 50,000 cells per well. Cells were maintained in DMEM (100 μl) media supplemented with 10% FBS, 100U/ml penicillin and 100 μg/ml streptomycin in a humidified atmosphere at 37°C in 5% CO_2_ for a day before the DMR experiment. Prior to proceeding with the DMR experiment, cells were washed with 1X HBSS buffer with HEPES (20 mM, pH 7.4) and allowed to temperature equilibrate in the same buffer (100 μl) for 1 h at room temperature. Basal DMR responses were monitored in Corning Epic BT system (Corning Epic-product code 5053) for 15 min to obtain a baseline reading that ensures the signal is stable and defined as zero. Thereafter, different concentrations of the compound of interest, AngII (final concentrations: 10 μM, 1 μM and 0.01 μM), were added and the DMR signal (in picometer) was monitored for 60-90 min. In case of allosteric modulator studies, before addition of the AngII, negative allosteric modulator DCP1-3 (final concentration 1 μM) and DCP1-16 (final concentration 0.1 μM) were incubated with cells in 1X HBSS buffer with HEPES (20 mM, pH 7.4) (100 μl) for 30 min^54^. Raw data was collected and analyzed in Corning-Epic Analyzer software to generate DMR response graphs (in picometer) over time (in minutes).

### PRESTO-TANGO β-arrestin recruitment assay

The assay was performed as previously described^32, 55–56^. HEK293 cell line stably expressing a tTA-dependent luciferase reporter and a β-arrestin2-TEV fusion gene (HTLA) cells (50,000/well) were seeded in a 6-well plate and transfected with 750 ng of each of the AT1R-TANGO plasmids using FUGENE 6 Transfection (PROMEGA). The following day, transfected HTLA cells (50,000 cells/well) were plated onto poly-L-lysine pre-coated 96-well microplates and allowed to attach to the plate surface for at least 12 hours prior to treatment. Cells were treated with ligands (AngII, DCP1-3, DCP1-16) and incubated overnight at 37°C, 5% CO_2_ in a humidified environment. The next day a volume of Dual-Glo® Reagent was added to each well, equal to the volume of culture medium in the well and mixed (75 μl of Reagent to cells grown in 75 μl of medium). 10 minutes were allowed for cell lysis to occur, and then the firefly luminescence was measured in a luminometer (Flexstation 2, Molecular devices). On a TriLux (Perkin-Elmer) plate counter. Data were normalized and analysed using the sigmoidal dose–response function built into GraphPad Prism 8.0.

### Reactive Oxygen Species (ROS)

To assess ROS levels, cells were plated and next day changed to serum free, phenol red free medium for 2 hours prior to treatment for 4 hours with AngII and incubated at 37°C 5% CO_2_. Cells were washed then incubated in 8 μM of the ROS sensitive probe 5-(and-6)-chloromethyl-2′,7′-dichlorodihydrofluorescein diacetate acetyl ester (CM-H2DCFDA) (Molecular Probes, Eugene Oregon), for 45 minutes. CM-H2DCFDA is a membrane-permeable compound that, upon internalization, is converted by intracellular esterases to the membrane impermeable compound, 2’ 7’ -dichlorofluorescein. In the presence of H_2_O_2_, this is compound is converted to the fluorescent compound, DCF. CM-H2DCFDA dye was replaced with serum free phenol red free medium prior to quantification. Data was quantified by measuring fluorescence intensity at 495 nm excitation and 517 nm emission (Flexstation 2, Molecular devices). Ter-butyl hydroperoxide was used as a positive control. Fluorescence values for AngII, AngII + Aab and AngII + Aab + DCP1-3 or DCP1-16 were compared using unpaired t-test with Welch’s correction in Graphpad Prism 8.

## Acknowledgements

We thank the LRI FACS and Protein and Peptide Biochemistry Cores and Dr. Stanely Hazen for providing access to DMR instrument. We thank Abdo Boumitri for construction of AT1R-tango plasmid used in our TANGO assays. We extend thanks to Earl Poptic and the Hybridoma Core for assistance in production of rabbit IgG. We thank Michael Weiner of the LRI Computer Support Core facility for Linux support. This work was supported by National Institutes of Health RO1 grants, HL132351 (SSK), HL142091 (SSK) and an Innovation Award from Lerner Research Institute.

Correspondence and requests for materials should be addressed to SSK.

## Author Contributions

DSK, carried out all computational studies, HTVS, characterization of allosteric ligands, signaling assays, myograph data acquisition, planning, analysis of data, preparation of figures and writing of the manuscript. ZPJ, standardization of myograph assay system, writing of IACUC protocol, mouse surgery, ex-vivo vasoconstriction data supervision, manuscript preparation. TH, assisted in ROS assay, myograph assay, manuscript preparation. PS, assisted in DMR assays. TP, structural analysis of small molecule hits, preparation of manuscript. RD, radio ligand binding assays, ELISA assays, molecular biology, manuscript preparation. SSK, procured funding, supervised data analysis, manuscript planning and managed the overall project. The manuscript was written by SSK and DSK, and included contributions from all authors.

---

**Supplementary Table 1.**
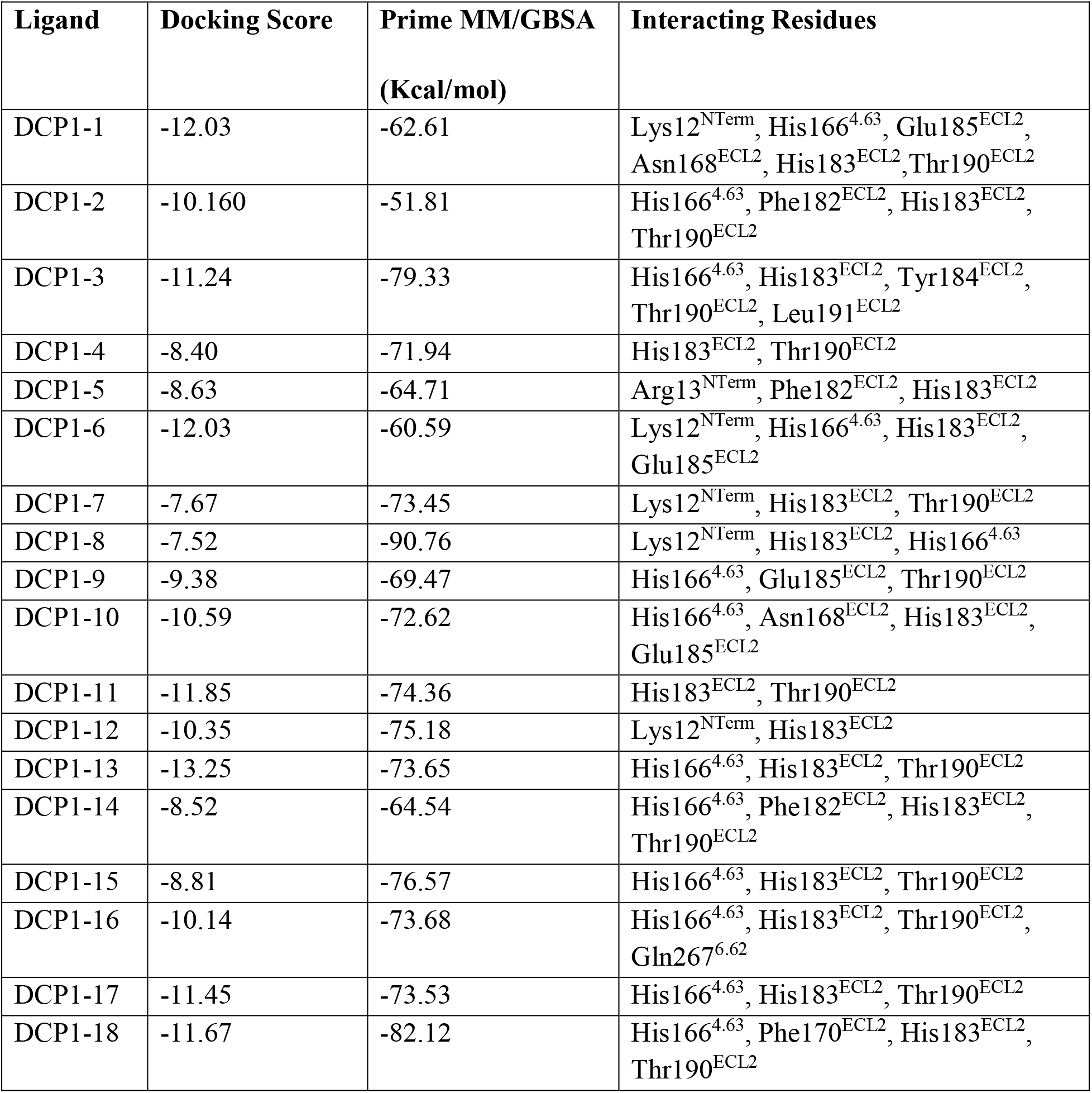
Docking score and Prime MM/GBSA free energy of all 18 allosteric hits (DCP1-compounds) from Induced Fit Docking (IFD) Pose with their interacting residues.

| Ligand | Docking Score | Prime MM/GBSA<br>(Kcal/mol) | Interacting Residues |
| --- | --- | --- | --- |
| DCP1-1 | -12.03 | -62.61 | Lys12<sup>NTerm</sup>, His166<sup>4.63</sup>, Glu185<sup>ECL2</sup>,<br>Asn168<sup>ECL2</sup>, His183<sup>ECL2</sup>, Thr190<sup>ECL2</sup> |
| DCP1-2 | -10.160 | -51.81 | His166<sup>4.63</sup>, Phe182<sup>ECL2</sup>, His183<sup>ECL2</sup>,<br>Thr190<sup>ECL2</sup> |
| DCP1-3 | -11.24 | -79.33 | His166<sup>4.63</sup>, His183<sup>ECL2</sup>, Tyr184<sup>ECL2</sup>,<br>Thr190<sup>ECL2</sup>, Leu191<sup>ECL2</sup> |
| DCP1-4 | -8.40 | -71.94 | His183<sup>ECL2</sup>, Thr190<sup>ECL2</sup> |
| DCP1-5 | -8.63 | -64.71 | Arg13<sup>NTerm</sup>, Phe182<sup>ECL2</sup>, His183<sup>ECL2</sup> |
| DCP1-6 | -12.03 | -60.59 | Lys12<sup>NTerm</sup>, His166<sup>4.63</sup>, His183<sup>ECL2</sup>,<br>Glu185<sup>ECL2</sup> |
| DCP1-7 | -7.67 | -73.45 | Lys12<sup>NTerm</sup>, His183<sup>ECL2</sup>, Thr190<sup>ECL2</sup> |
| DCP1-8 | -7.52 | -90.76 | Lys12<sup>NTerm</sup>, His183<sup>ECL2</sup>, His166<sup>4.63</sup> |
| DCP1-9 | -9.38 | -69.47 | His166<sup>4.63</sup>, Glu185<sup>ECL2</sup>, Thr190<sup>ECL2</sup> |
| DCP1-10 | -10.59 | -72.62 | His166<sup>4.63</sup>, Asn168<sup>ECL2</sup>, His183<sup>ECL2</sup>,<br>Glu185<sup>ECL2</sup> |
| DCP1-11 | -11.85 | -74.36 | His183<sup>ECL2</sup>, Thr190<sup>ECL2</sup> |
| DCP1-12 | -10.35 | -75.18 | Lys12<sup>NTerm</sup>, His183<sup>ECL2</sup> |
| DCP1-13 | -13.25 | -73.65 | His166<sup>4.63</sup>, His183<sup>ECL2</sup>, Thr190<sup>ECL2</sup> |
| DCP1-14 | -8.52 | -64.54 | His166<sup>4.63</sup>, Phe182<sup>ECL2</sup>, His183<sup>ECL2</sup>,<br>Thr190<sup>ECL2</sup> |
| DCP1-15 | -8.81 | -76.57 | His166<sup>4.63</sup>, His183<sup>ECL2</sup>, Thr190<sup>ECL2</sup> |
| DCP1-16 | -10.14 | -73.68 | His166<sup>4.63</sup>, His183<sup>ECL2</sup>, Thr190<sup>ECL2</sup>,<br>Gln267<sup>6.62</sup> |
| DCP1-17 | -11.45 | -73.53 | His166<sup>4.63</sup>, His183<sup>ECL2</sup>, Thr190<sup>ECL2</sup> |
| DCP1-18 | -11.67 | -82.12 | His166<sup>4.63</sup>, Phe170<sup>ECL2</sup>, His183<sup>ECL2</sup>,<br>Thr190<sup>ECL2</sup> |

**Supplementary Table 2.** Prime MM/GBSA free energy of binding sAngII to AT1R in the absence and presence of DCP1-3 and DCP1-16. Prime MM/GBSA binding free energy was calculated from the final pose of 1000 ns MD simulation. MD simulations were run in triplicate. As a hallmark of an allosteric compound, a decrease in the binding affinity of the orthosteric ligand sAngII was evident upon binding of DCP1-3 and DCP1-16. The size of the binding site of G-protein decreased upon binding of DCP1-3. In the case of DCP1-16, interestingly, reduction in the size of the cytoplasmic cleft compared to the sAngII bound AT_1_R structure was not observed. This may be the basis for β-arrestin bias property exhibited by of DCP1-16 in Fig 3e.

| Ligand | Prime MM/GBSA (Kcal/mol) |
| --- | --- |
| sAngII | -148.53 ± 6.48 |
| sAngII (+DCP1-3) | -139.42 ± 22.29 |
| sAngII (+DCP1-16) | -138.96 ± 4.69 |

**Supplementary Fig. 1.**
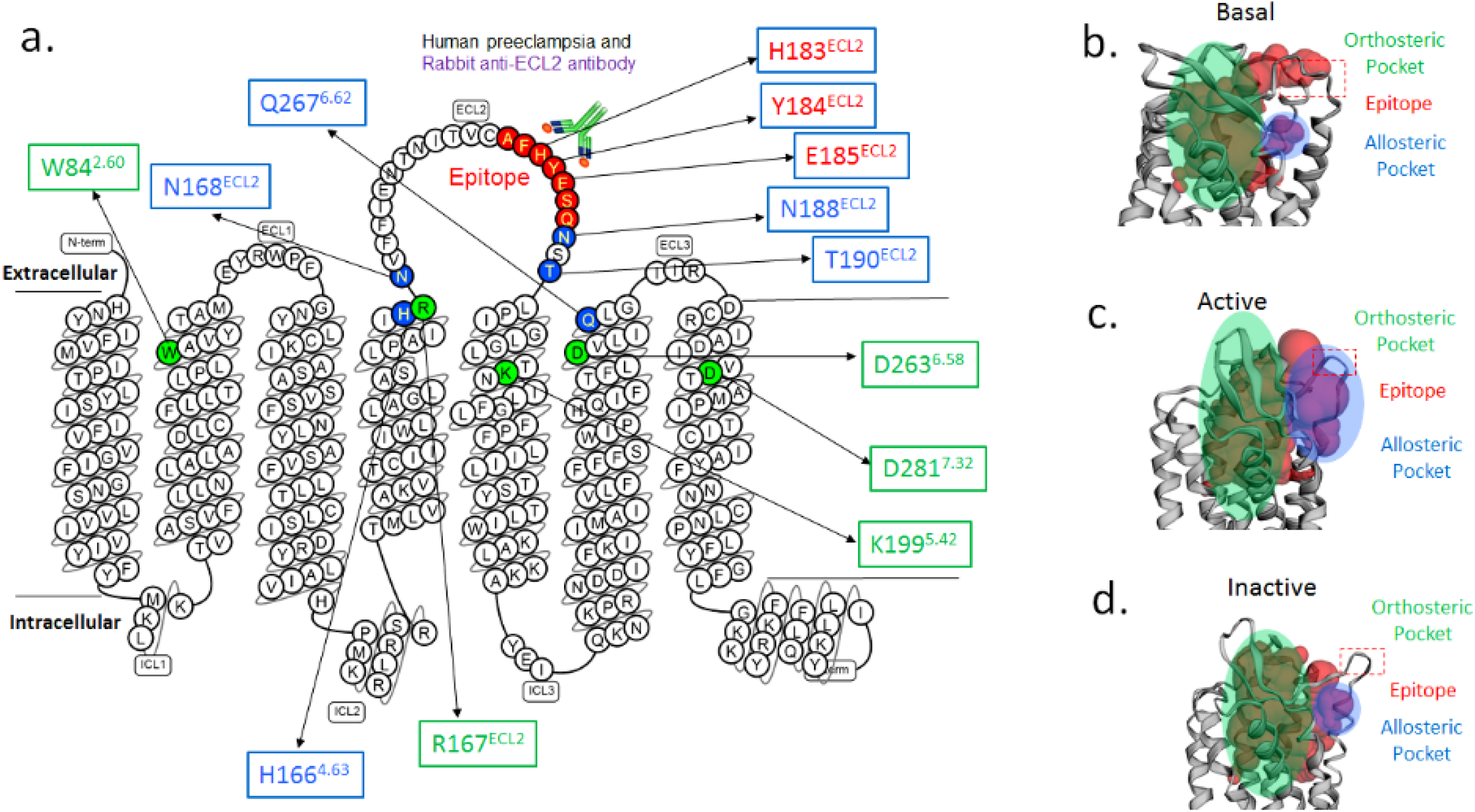
AT1R structure and shapes of cryptic allosteric pocket in different states. **a,** Secondary structure map with highlighted preeclampsia epitope sequence (red), residues constituting the allosteric pocket (blue), orthosteric pocket (green) and autoantibody binding site (red). **b-d**, Allosteric pocket (purple shade) predicted by CASTp^27^ is distinct from AngII binding pocket in active state AT1R. Change in the size and shape of allosteric pocket shown in basal, active and inactive states.

**Supplementary Fig. 2.**
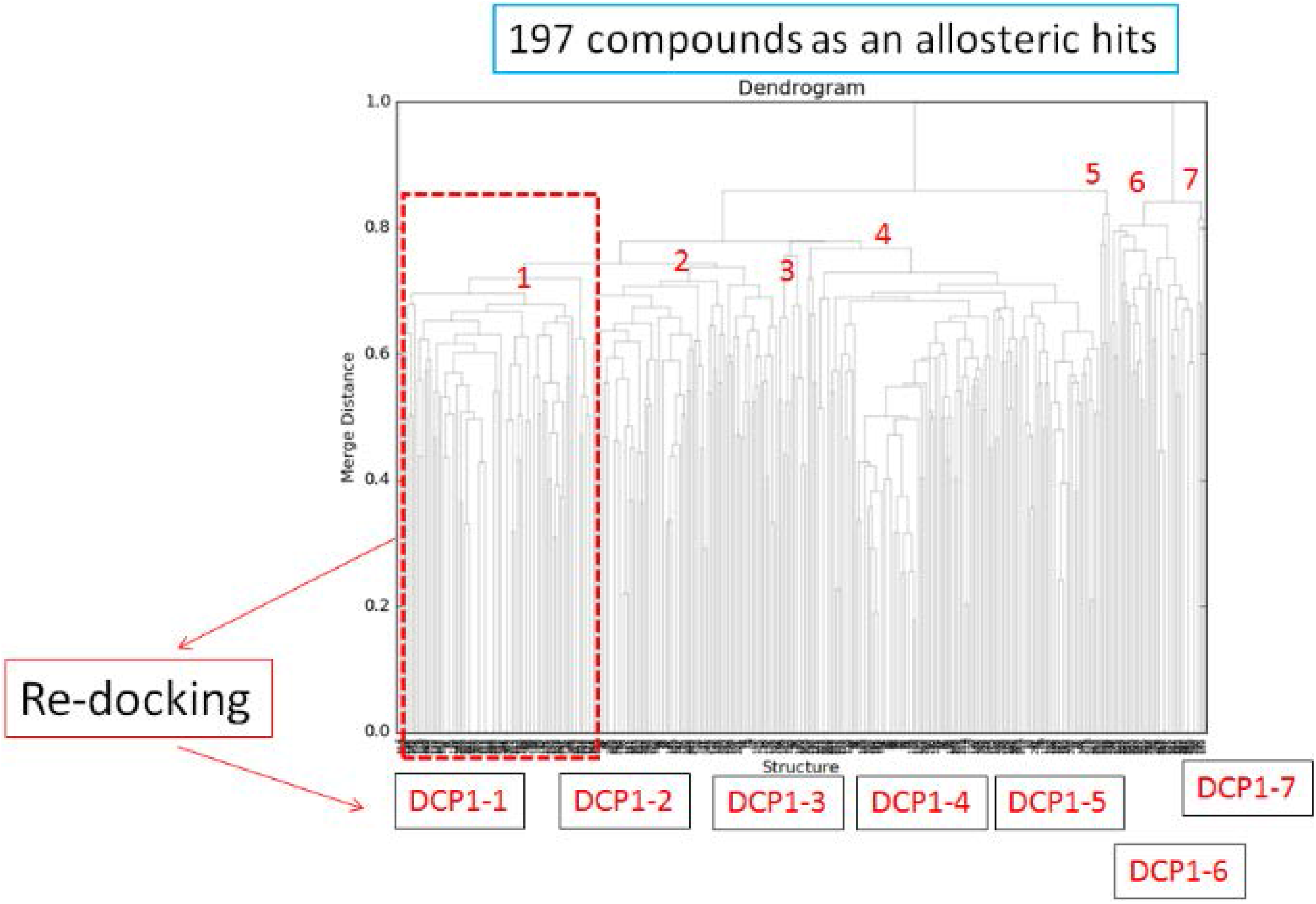
Clustering analysis of 197 hits. Compounds are clustered based on their structural similarity. Compounds from each cluster were re-docked and the compound with best score was selected and code-named as DCP1-1 to DCP1-7.

**Supplementary Fig. 3.**
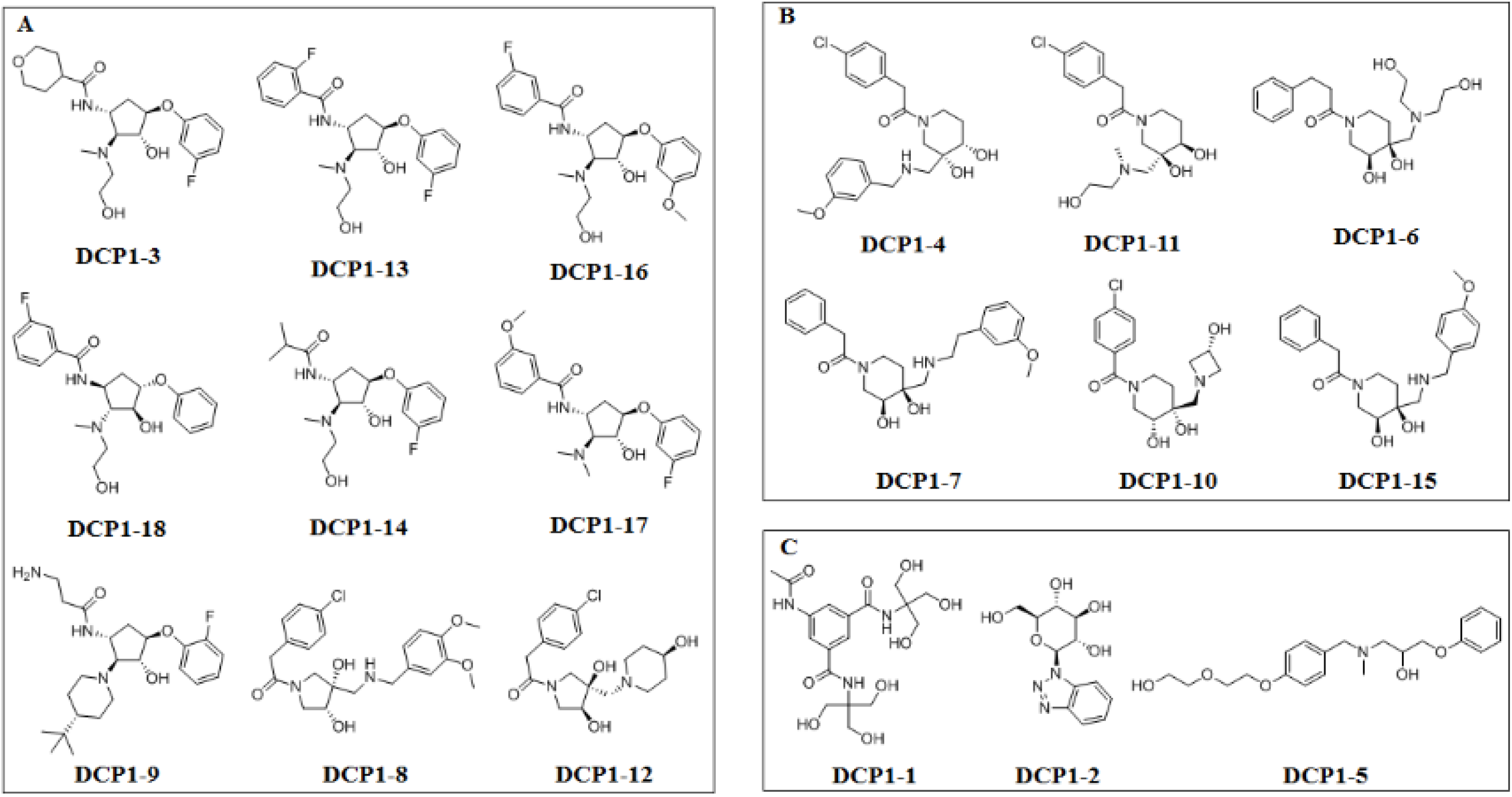
Structures of compounds DCP1-1 to DCP1-18 identified as AT_1_R allosteric site binders. Based on their structures, the18 compounds fall into 3 classes: (A) Cyclopentane and Pyrrolidine derivatives; (B) Piperidine derivatives; (C) Distinct structures. Note: compounds names are in the order of discovery and interacting residues for all are provided in Supplementary Table 1.

**Supplementary Fig. 4.**
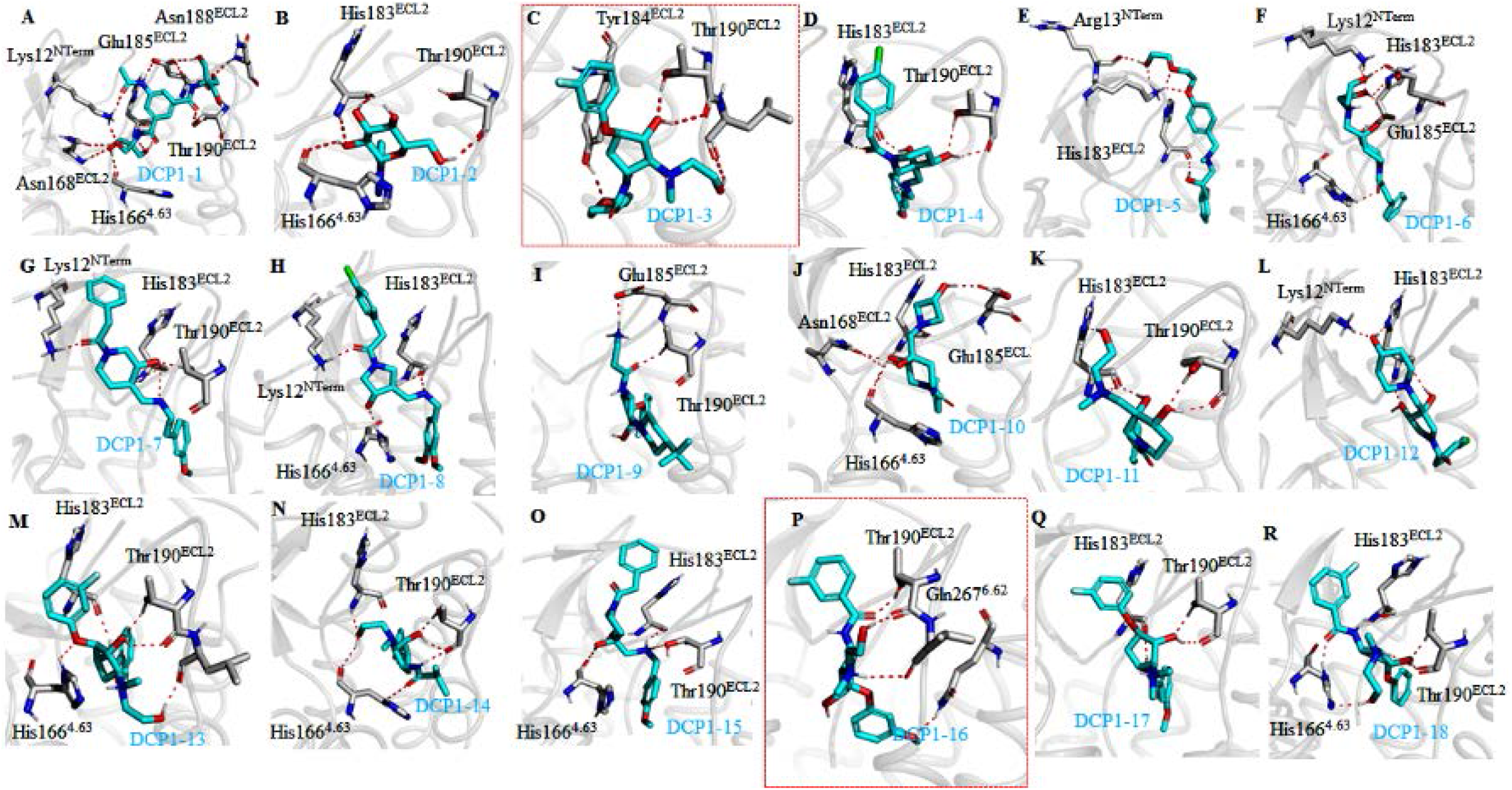
Binding poses of DCP1-1 to DCP1-18. Note that two compounds of interest in our study, DCP1-3 and DCP-16 have four functional groups attached to the cyclopentane core. Comparison of DCP1-3 and DCP-16 poses indicates that residues, His166^ECL2^, His183^ECL2^, Tyr184^ECL2^, Thr190^ECL2^ and Leu191^ECL2^ participate in binding. Both contain the same functional groups at position 2 (N-methylethanolamine hydroxyl forming ionic bond with His166^4.63^ and H-bond with carboxylic group of Leu191^ECL2^) and at position 3 (the hydroxyl forming a strong H-bond with the carboxylic group of Thr190^ECL2^). The tetrahydropyran-4-carboxamide group at position 1 in DCP1-3 forms an H-bond with hydroxyl of Tyr184^ECL2^. The position 4 aromatic ring (benzene-3-fluorophenol) is involved in a π-π interaction with His183^ECL2^ in DCP1-3. In DCP1-16 the carbonyl fluorobenzamide at position 1 is engaged in H-bond interaction with Thr190^ECL2^. The position 4 methoxyphenol group in DCP1-16, forms a hydrogen bond with Gln267^6.62^ and is projected towards the orthosteric site. Majority of the functional groups in DCP1-16 interact with AT1R, which might explain why DCP1-16 is more potent than DCP1-3. An interesting observation made is that fluorobenzene and ethanol attached in two functional groups do not contribute substantially to interaction with the receptor. Note: also see Supplementary Table 2. These observations will help us in further designing novel, more potent allosteric leads for AT1R.

**Supplementary Fig. 5.**
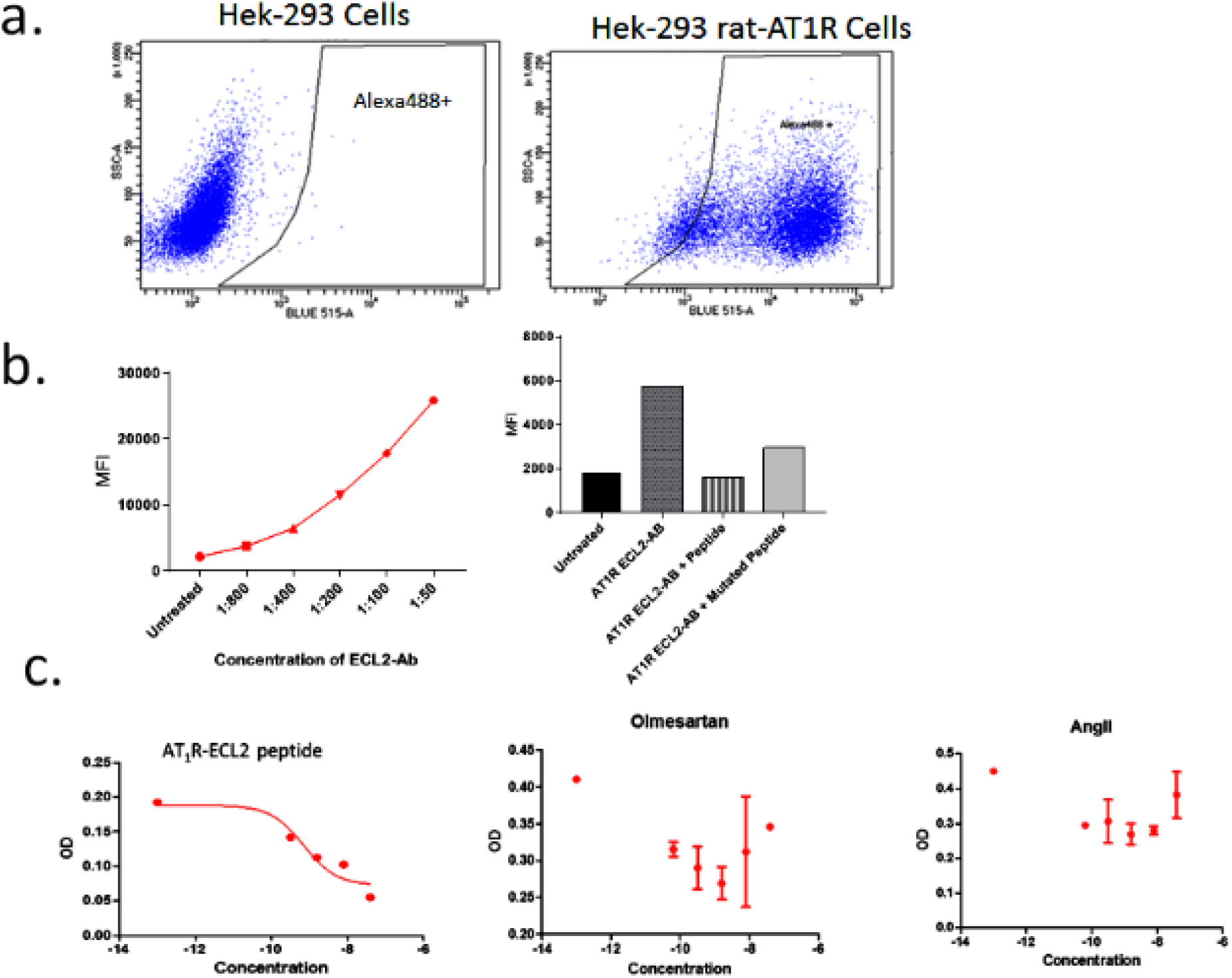
Generation and characterization of rabbit IgG for the preeclampsia epitope of AT1R: Polyclonal antibodies were raised by immunizing rabbits with ECL2 peptide hapten, –CIENTNITVS<u>AFHYESQNS</u>–. IgG purified from immunized rabbit serum by epitope peptide affinity chromatography were characterize as shown: **a**, In Fluorescence-activated cell sorting (FACS) experiment, the IgG bound to HEK-AT1R cells but not to HEK293 cells. **b**, The mean fluorescence intensity (MFI) signal changed in proportion to antibody dilution (left panel). The bar-graphs in right panel show that MFI signal of IgG treated HEK-AT1R was inhibited by the preeclampsia epitope-peptide, –AFHYESQNST–. Blocking efficacy was reduced when we mutated one amino acid in the epitope to the sequence –AFH**<u>A</u>**ESQNST–. This result suggests that the antibody is highly specific to the AT1R’s preeclampsia epitope. **c**, IgG binding to HEK-AT1R was blocked by the antigenic peptide, but was not inhibited by orthosteric ligands, AngII and Olmesartan.

**Supplementary Fig. 6.**
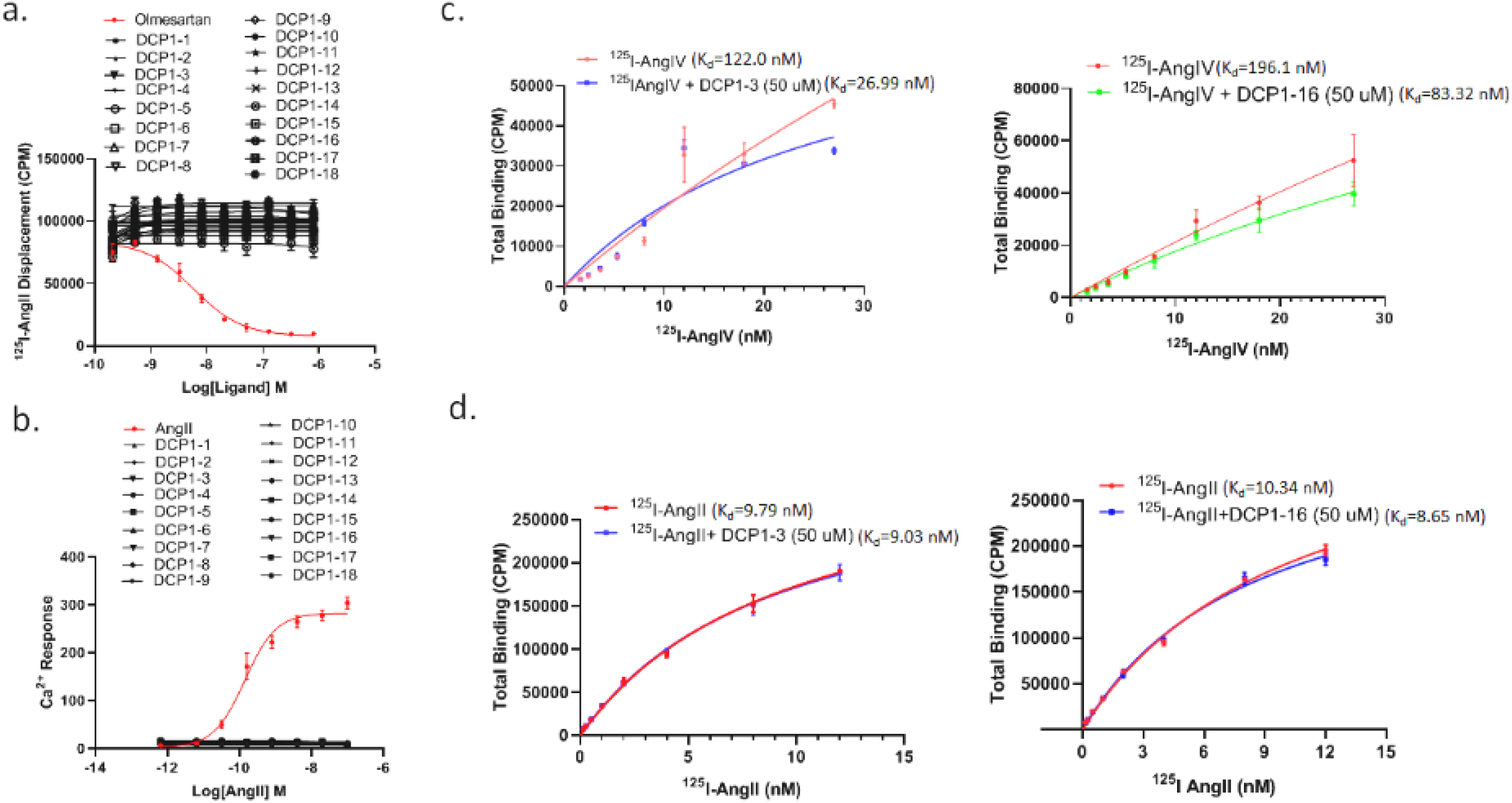
Effect of allosteric hits on binding of orthosteric ligand and calcium signaling: **a,** Allosteric ligands did not affect the binding of orthosteric ligand ^125^I-[Sar^1^,Ile^8^]AngII in HEK-AT1R membranes, while the antagonist Olmesartan did as expected. **b**, To assess potential ago-PAM property, treatment of HEK-AT1R cells with 18 allosteric compounds did not induce calcium mobilization while the agonist AngII did as expected. **c,** Effect of 50 μM DCP1-3 and DCP1-16 on binding of orthosteric agonist, ^125^I-AngIV. **d**, Effect on ^125^I-AngII binding.

**Supplementary Fig. 7.**
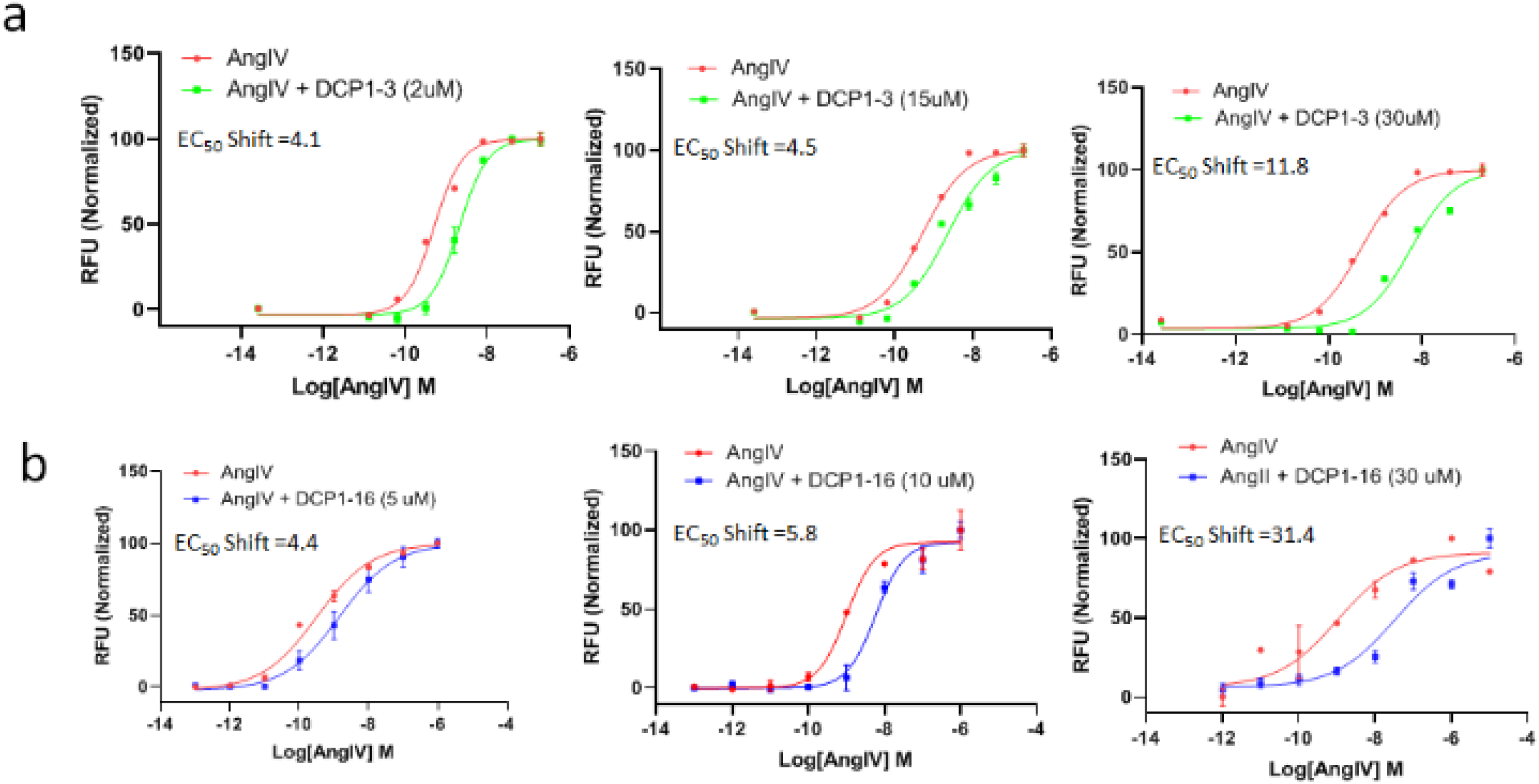
Efficacy modulation by DCP1-3 and DCP1-16. **a,** Fold-shift of AngIV EC_50_ induced by concentrations of DCP1-3 indicated. **b**, Fold-shift of AngIV EC_50_ induced by concentrations of DCP1-16 indicated.

**Supplementary Fig. 8.**
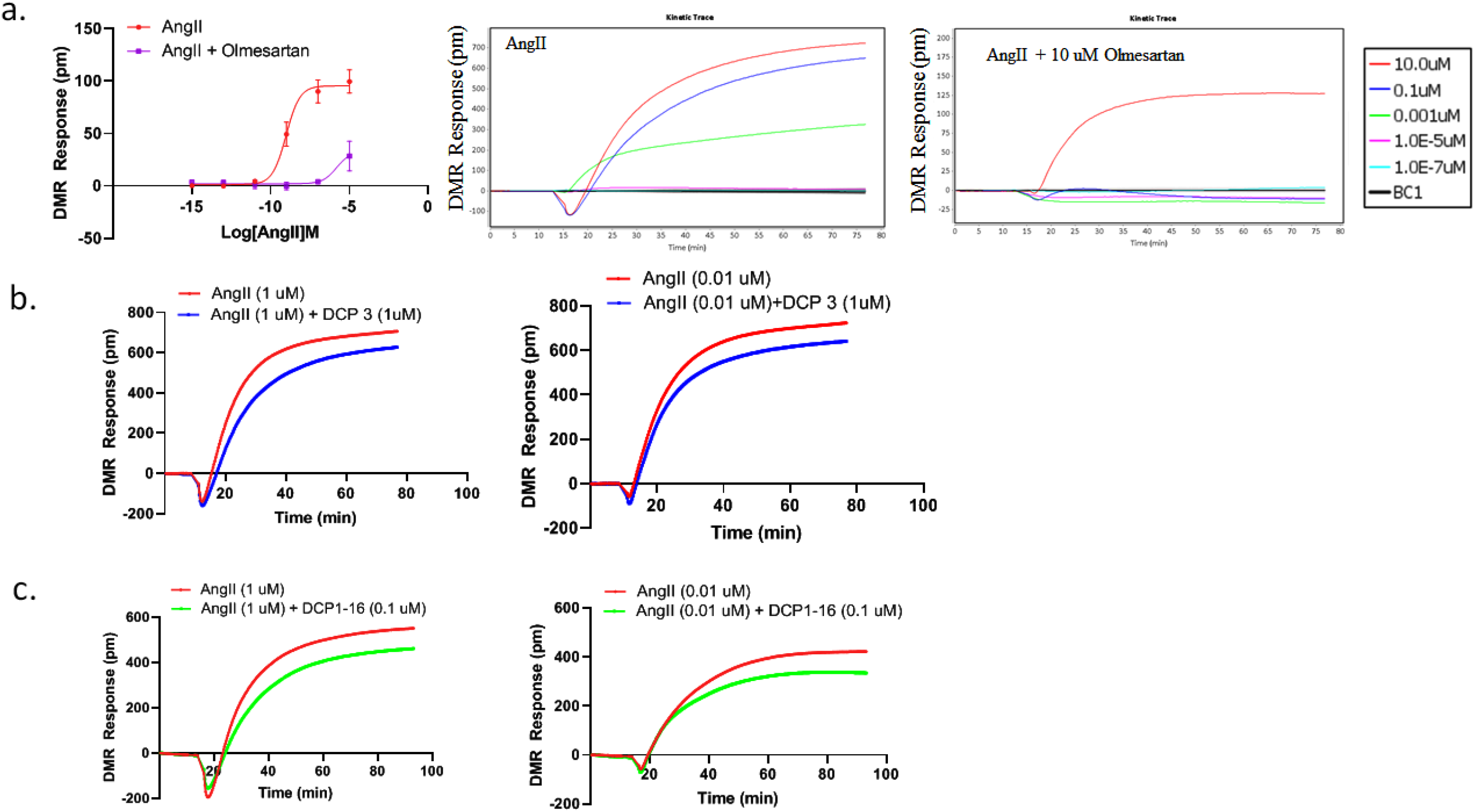
DMR response modulation by DCP1-3 and DCP1-16. **a,** Concentration-response curves. In the left panel each data point represents average ± S.E. from three independent experiments. Right panels show kinetic traces for single concentrations of ligands, which is used to calculate the full dose response curves. **b**, Effect of DCP1-3 on kinetic traces of indicated concentrations of AngII. **c**, Effect of DCP1-16 on kinetic traces of indicated concentrations of AngII. The single concentration kinetic trace data was used to represent the dose-response curves presented in Fig. 3c. *BC1, Buffer control.

**Supplementary Fig. 9.**
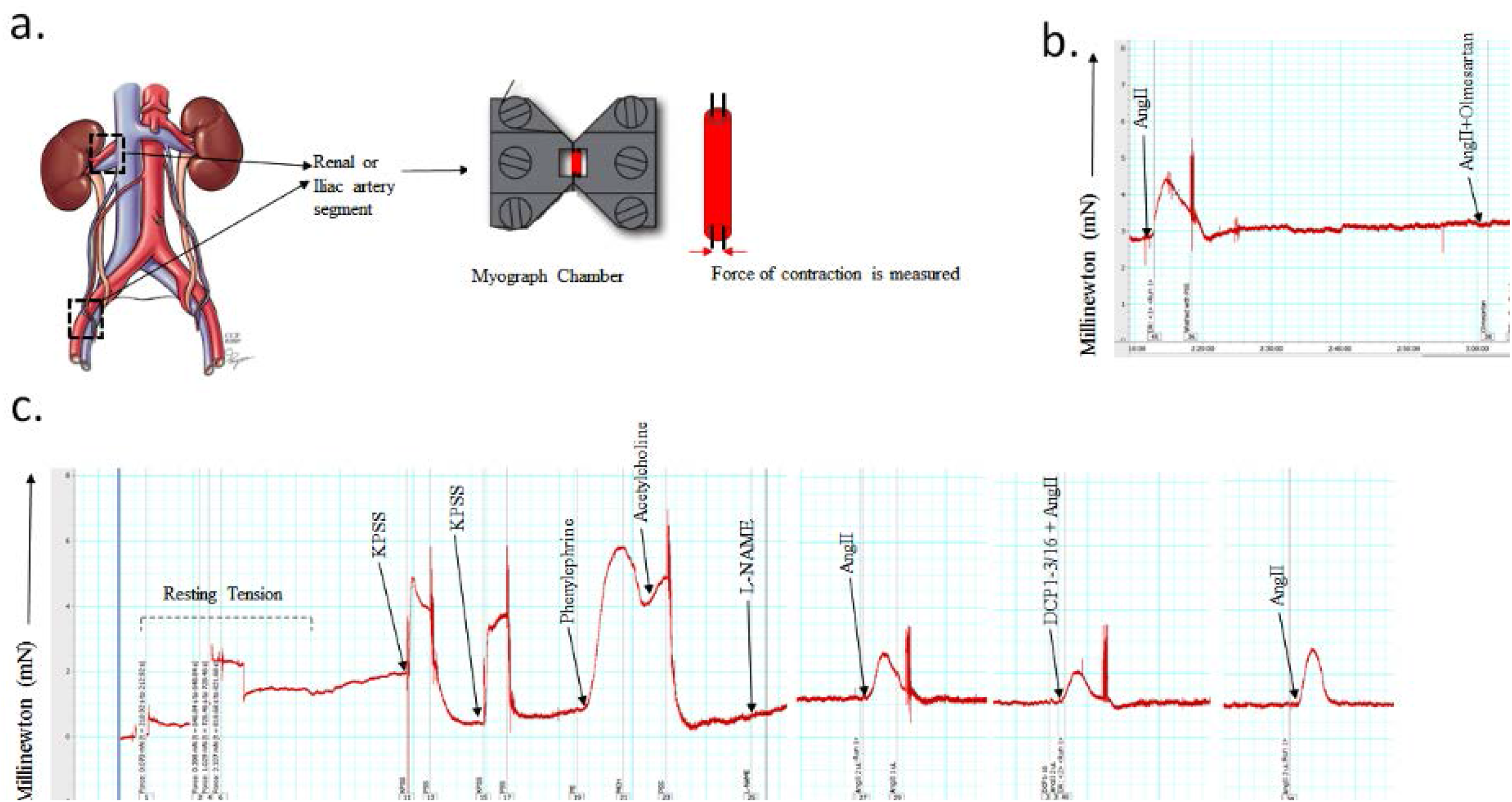
Myograph measurement of vasoconstriction response. **a,** Representation of renal and iliac arterial segment explants mounted into tension-wires in the myograph chamber. **b**, A representative trace of constriction response produced upon treatment with AT1R agonist and antagonist ligands. **c**, A representative response-trace is shown along with calibration of basal tension, assessment of transient response of the mounted-explant to sequential treatment starting with KPSS, followed by pharmacological ligands indicated. Area under the peak for each treatment condition is used to represent average ± S.D. in Fig. 3D.

